# A structure and function-based complete mutational map of Human Hemoglobin using AI

**DOI:** 10.1101/2025.09.30.679504

**Authors:** Franco Salvatore, Franco G. Brunello, Claudio D. Schuster, Marcelo A. Martí

## Abstract

Hemoglob+in (Hb), a well-characterized protein central to oxygen transport and molecular medicine, serves as a model for studying how sequence variations influence protein structure and function. Its precise activity depends on tightly regulated structural dynamics, which can be disrupted by mutations that give rise to structural hemoglobinopathies—including sickle cell disease, unstable hemoglobins, methemoglobins, and hemoglobins with altered oxygen affinity—each associated with distinct functional and clinical consequences.Among genetic variants, missense mutations are the most widely studied in clinical settings. Accurately predicting their clinical impact remains challenging, requiring integration of evolutionary, biochemical, and structural data. While broad deep learning models like AlphaMissense show promise, they often lack interpretability and protein-specific precision. This motivates the development of focused models that leverage detailed knowledge of individual proteins, like hemoglobin, to improve both predictive power and mechanistic understanding.

In this work, we conducted a comprehensive analysis of all known and potential human adult hemoglobin (HbA) variants, guided by the hypothesis that a deep understanding of the sequence–structure–function relationship in Hb can yield interpretable and predictive insights into the functional and clinical consequences of single amino acid substitutions. We curated an updated dataset of HbA variants annotated with their clinical classifications—Benign, Pathogenic, or of Uncertain Significance (VUS)—and systematically mapped each to a range of features, including structural location and classification, predicted impact on folding stability, and evolutionary conservation. Using this data, we developed a pathogenicity prediction model and benchmarked it against AlphaMissense, demonstrating strong and complementary performance. Additionally, we generated a complete mutational landscape of all possible single amino acid substitutions (SAS) in HbA, providing a resource for future clinical interpretation.

Our findings provide insight into the molecular basis for variant effects in HbA and highlight the utility of combining structure-informed features with Machine Learning (ML) for variant interpretation. Moreover, our results offer a framework for evaluating the portability and interpretability of variant effect predictors across structurally dynamic systems, with implications in the improvement of variant classification in other protein families.

## Introduction

Hemoglobin (Hb), a tetrameric protein composed of two α and two β globin chains, is one of the most extensively studied proteins in humans, both from a clinical and structural biology perspective [10–15]. Its critical role in oxygen transport, and its involvement in some of the first genetic diseases ever described, such as sickle cell disease and the thalassemias, have made Hb a cornerstone model in molecular medicine. Beyond its biomedical relevance, Hb offers a uniquely well-characterized system for exploring the relationship between protein sequence, three-dimensional structure, and functional outcome. Moreover, high-resolution crystal structures of Hb in both its oxygenated (R) and deoxygenated (T) states, also allow for an in-depth examination of how conformational dynamics and allosterism influence protein function and, consequently, how genetic variants may disrupt it.

The α- and β-globin chains of HbA are globular domains organized in 8 α-helices (named A to H) that surround the heme group. The central iron atom of the heme is coordinated on its proximal side by the conserved HisF8 residue. Hemoglobin’s function, oxygen transport, relies on the reversible binding of oxygen to the ferrous heme iron within the distal cavity. Its affinity is tightly regulated by key residues that allow oxygen entry and release, as well as its stabilization upon coordination, such as the distal HisE7 and ValE11. Furthermore, a key aspect to HbA function is the allosteric modulation of oxygen affinity, achieved by switching between the low-affinity Tense (T) state and the high-affinity Relaxed (R) state. This transition is physiologically modulated by factors such as pH (the Bohr effect), temperature, the presence of allosteric ligands such as 2,3-bisphosphoglycerate (2,3-BPG). Additionally, oxygen affinity can be altered by changes in the oxidation state of the heme iron. All these regulations depend on a tight modulation of Hb structure and dynamics, which is encoded in the underlying subunit sequences [10]. Therefore, a deep understanding of the sequence–structure–function relationship is essential to explain how mutations can alter oxygen affinity, disrupt folding, or promote aggregation, ultimately leading to hemoglobinopathies.

Hemoglobinopathies are a group of inherited disorders caused by structural or regulatory mutations in the Hb molecule, affecting its function, stability, or expression. These conditions are broadly classified into two categories: structural variants and synthesis disorders (e.g., thalassemias). Structural mutations may alter hemoglobin’s stability, oxygen affinity, T-to-R allosteric transitions, or its capacity to form stable tetramers. A well-known example is sickle cell disease (HbS), caused by a p.Glu6Val substitution in the β-globin chain [16]. This mutation promotes the polymerization of deoxygenated Hb, forming rigid fibers that distort red blood cells into a sickle shape. These misshapen erythrocytes have reduced lifespan and tend to obstruct capillaries, leading to vaso-occlusive crises, hemolysis, and progressive organ damage. Other clinically relevant variants include high-affinity hemoglobins, which impair oxygen release to tissues and lead to compensatory erythrocytosis, and low-affinity variants, which can cause peripheral cyanosis due to elevated levels of deoxygenated hemoglobin in arterial blood, despite preserved tissue oxygenation. Unstable hemoglobins may precipitate within erythrocytes, forming Heinz bodies and leading to hemolytic anemia. In contrast, variants that promote methemoglobinemia (met-Hb) -characterized by the accumulation of ferric (Fe^3+^) Hb, which is unable to bind oxygen and simultaneously increases the oxygen affinity of the remaining ferrous subunits-result in functional hypoxia and cyanosis. Together, these diverse clinical manifestations underscore the critical impact of specific genetic mutations on hemoglobin structure and function.

Missense variants, which result in the substitution of one amino acid for another in a protein sequence -commonly referred to as Single Amino Acid Substitutions (SAS)-are among the most prevalent forms of genetic variation in humans. While many of these variants are classified as Benign (B) due to their lack of impact on health, others can severely alter protein function, leading to Mendelian diseases and thus being classified as Pathogenic (P). Predicting the pathogenicity of missense variants remains a central challenge in human genetics, largely due to the complexity and diversity of the underlying molecular mechanisms. Although advances in high-throughput sequencing have significantly increased our ability to detect such variants, accurate interpretation requires the integration of multiple layers of biological data, including evolutionary conservation, physicochemical context, and structural constraints.

Over the past two decades, numerous computational tools have been developed to predict the pathogenicity of missense variants, using diverse biological signals. Early approaches, such as SIFT and PolyPhen-2, relied on evolutionary conservation, amino acid properties, and, when available, structural features to estimate the impact of SAS. Others, like PROVEAN and LRT, expanded on this by incorporating pairwise alignment strategies or codon-level selection models. Additional layers of biological information were integrated in tools like fitCons, which leveraged functional genomic data, or CADD, which used simulated data to extend predictions genome-wide. More recently, ensemble models such as REVEL have emerged, combining outputs from multiple predictors using machine learning to improve accuracy and robustness. This evolution reflects a shift from conservation-based heuristics to data-driven models that integrate sequence, structure, and functional context, offering increasingly reliable variant effect predictions.

One notable example is AlphaMissense (AM) [2], a deep learning model developed by DeepMind (a Google company) to predict the pathogenicity of all possible missense variants in the human proteome. The model builds on the architecture and internal representations of AlphaFold, leveraging protein sequence embeddings that capture patterns from large-scale evolutionary and structural data. Notably, AM does not incorporate explicit 3D structural inputs; instead, it relies on sequence-derived features processed through a neural network trained on clinically annotated variants to generate pathogenicity scores. While AM demonstrates strong performance on genome-wide benchmarks, it remains a black-box model whose predictions offer limited mechanistic interpretability. Moreover, because it is designed to generalize across the entire human proteome, it may overlook protein-specific constraints or functional nuances, particularly in well-characterized systems such as hemoglobin. This limitation motivates our study: a model trained on data from a single protein or protein family, enriched with detailed structural and functional knowledge, can not only outperform broad-spectrum models like AM in specific domains, but also yield more interpretable and biologically grounded insights. This highlights a central question in molecular biology: to what extent can the sequence-structure-function paradigm be leveraged not only to explain, but also to predict, the clinical impact of missense variants?

In the present work, we conducted an exhaustive study of known and potential Hb genetic variants, under the hypothesis that a deep understanding of sequence-structure-function relationships in Hb can yield interpretable and predictive insights into the functional and clinical impact of SAS. We compiled and curated an up-to-date dataset of Hb variants with known clinical effect, and mapped each SAS to multiple features, including topological location, structural classification, folding stability, and evolutionary conservation. Using this annotated dataset, we developed a predictive model of variant pathogenicity and evaluated its performance in comparison to, and in combination with, AlphaMissense, showing an excellent performance. Finally, we provide a complete mutational map of all possible SAS in HbA, intended to support future clinical interpretation and decision-making. Ultimately, this work aims to bridge the gap between mechanistic insight and predictive modeling in variant interpretation, illustrating how structural biology can inform and enhance the evaluation of genetic variation in clinically important proteins such as HbA.

## Computational Methods

### Hb Sequence and Structure Data: Download and Processing

Canonical sequences for the HBA and HBB protein subunits were retrieved from UniprotKB [17] in FASTA format using the following UniProt IDs: P69905 (HBA) and P68871 (HBB). Structural data corresponding to the oxy (R-state) and deoxy (T-state) conformations of tetrameric hemoglobin A (HbA) were downloaded from the Protein Data Bank (PDB) [18] under the accession codes 1GZX (oxy) and 2DN2 (deoxy). Pairwise sequence alignments were performed to map the residues from the PDB structures to the canonical UniProt numbering, which is consistently used throughout the work.

Using the aforementioned structures, all HbA residues were structurally classified into one of five structural categories: i) Surface ii) Interface iii) Active Site iv) Core, or v) Switch residues. Surface residues were assigned based on residue depth (RD) [19], calculated as the shortest distance from the residue’s center of mass (COM) to the molecular surface. In the case of glycines, the Cα atom was used as a proxy for the COM. If the COM was closer to the surface than the residue Cα atom, the residue was classified as exposed; otherwise, it was considered buried. The surface is defined by those atoms that define the corresponding solvent-accessible surface area (SASA). Other strategies to define surface residues, based on residue SASA or relative distance to the protein’s center of mass (normalized by the radius of gyration) were evaluated, but RD yielded the most consistent classification. Interface residues were defined as those whose COM lay within 8Å of the COM of any residue from a different subunit. Active site residues were defined as those located within 8Å of the heme iron atom. All remaining residues were classified as core. In cases where residues fulfilled multiple criteria, a hierarchical classification was applied: active site > interface > surface. Additionally, a small set of residues known to interact with 2,3-DPG (which binds to the central cavity of HbA, stabilizing its T state and reducing its affinity for oxygen) were annotated based on literature, though they were excluded from further analysis due to their limited number. Lastly, switch residues were defined as those whose structural classification changed between the T and R-states, reflecting conformational flexibility relevant to hemoglobin’s function.

For each residue, we also determined its secondary structure based on its phi/psi angles. Additionally, we computed the Jensen-Shannon conservation score [1], which quantifies the similarity between the amino acid distribution at a given position in a Multiple Sequence Alignment (MSA) and a reference distribution. Higher scores reflect stronger evolutionary conservation. To calculate conservation scores for HbA residues, we generated MSAs using MAFFT [20], based on 2905 mammalian hemoglobin sequences with a sequence length ranging from 100 to 150 amino acids, retrieved from UniProtKB [17].

### Variant properties download and assignment

Previously reported variants in the HBA1/HBA2 and HBB genes were retrieved mainly from the HbVar database [6] and were manually curated. The primary dataset from HbVar initially included 979 HbA variants, which were carefully reviewed and cross-validated with other databases such as ClinVar [7], gnomAD [8] and LOVD [9] to ensure consistency in clinical classification and, when possible, retrieve extra information of the variants. Each variant was assigned a pathogenicity label -(Likely) Pathogenic (P), Variant of Uncertain Significance (VUS), or (Likely) Benign (B)-. For most variants, pathogenicity annotations were consistent across databases, with only minor discrepancies that were manually resolved. Pathogenic variants from HbVar were further annotated with specific clinical phenotypes: i) methemoglobinemia (met-Hb), ii) altered oxygen affinity, and iii) Hb instability. Variant allele frequencies were obtained directly from gnomAD when available. After manual curation, the dataset was refined to 722 well-characterized variants (413 P and 309 B); the remainder (256 variants), classified as VUS or lacking clinical annotation, were excluded from model training.

For each known (or potential) variant, we computed changes in residue hydropathy (Δhyd), residue mass (Δmass), and substitution conservation using the BLOSUM62 matrix [21]. Additionally, we annotated the chemical properties of the wild-type and mutant amino acids. This feature, termed Chemical Mutation, categorizes amino acids based on their chemical nature; for example, a mutation from alanine to phenylalanine would be labeled as a hydrophobic-to-aromatic substitution.

### Determination of variant folding free energy change (ΔΔGfold) using FoldX

To assess the effect of each potential variant on protein stability, we used FoldX to compute the folding free energy change (ΔΔGfold) for every possible SAS at each residue of both HbA and HbB subunits, in both the oxy and deoxy structures, as described in previous studies [5]. Additionally, we computed the protein-protein interaction free energy change (ΔΔGint) using FoldX for residues located at the intersubunit contact surfaces.

### AlphaMissense

AlphaMissense (AM) [2] is a deep learning model developed by Google DeepMind to predict the pathogenicity of all possible missense variants across the human proteome. Building upon the architecture and internal representations of AlphaFold2, AM leverages protein sequence embeddings that capture evolutionary and structural patterns derived from large-scale sequence data. Notably, AM does not rely on explicit three-dimensional structural information; instead, it processes sequence-derived features through a neural network trained on a comprehensive set of clinically annotated variants to assign a pathogenicity score. This score ranges from 0 to 1, indicating the approximate probability of a given variant being clinically pathogenic. Based on thresholds defined in the original AM study, variants with scores below 0.34 are classified as likely benign, those above 0.56 as likely pathogenic, and intermediate values are considered variants of uncertain significance (VUS). All scores used in this work were obtained from a publicly available dataset released by DeepMind, which provides predictions for every possible missense mutation in the human proteome, in this case we retrieved the scores for all SAS mutations on HBA1/2 and HBB genes.

In the present study, AM was employed in two complementary ways: (i) as an independent benchmark model to assess the performance of our approach relative to an established state-of-the-art predictor; and (ii) as an additional input feature in a combined machine learning model (Combined Model, CM), alongside our curated features, to improve the prediction of clinically relevant variants in HbA.

### Machine Learning (ML) Model Building and Training

Several machine learning models were constructed and tested for their capacity to predict the pathogenicity of the collected HbA variants. The input features included: structural classification, simple topological notation (i.e. the unnumbered topological notation), chemical classification of native and alternative residues (i.e. Chemical Mutation), conservation score from mammalian hemoglobins, ΔHydropathy, ΔΔG FoldX stability, ΔΔG FoldX binding, BLOSUM62 substitution score, ΔMass and the AM probability score.

A 7-fold cross-validation strategy [22] was employed, with each fold containing approximately 103 variants. The entire procedure was repeated 50 times to reduce the risk of overfitting due to a fixed data split and enhance evaluation robustness. Each fold was balanced to maintain the original dataset distribution (∼57% pathogenic and ∼43% benign variants), thus improving model generalizability.

A range of canonical machine learning models were tested by global accuracy, including Random Forests [23], Support Vector Machines [24], and Xtreme Gradient Boosting [25]. Multiple feature combinations were evaluated following a logical selection scheme that avoided including redundant or highly correlated features within the same model. For instance, features describing similar properties—such as ΔHydropathy and ΔMass, or BLOSUM scores and Chemical Mutation—were not used together. Hyperparameters were optimized to improve classification performance. The final selected model was a Random Forest trained on the following features: structural classification, Simple Topological Notation, chemical classification of both native and mutant residues (i.e., Chemical Mutation), ΔΔG FoldX stability, ΔHydropathy, and conservation score.

To assess the contribution of AM, two versions of the model were constructed: one excluding the AM score (Model 1, M1), and one including it (Combined Model, CM). Both models were benchmarked against each other, and against AM alone, enabling direct evaluation of the added value of AM relative to the other curated features.

### M1 and CM pathogenicity thresholds

A key aspect of variant classification is determining optimal probability thresholds. Clinical significance predictions are based on a probability score indicating the likelihood that a variant belongs to the pathogenic class. Variants with scores above or below defined thresholds are assigned accordingly, while those falling in between are classified as VUS. To define optimal thresholds, we computed class-specific precision across a range of probability cutoffs (0–1). Final thresholds were selected to ensure a minimum precision of 85% for both pathogenic and benign classifications. The resulting VUS range corresponds to the interval between these two thresholds, where variant scores do not meet the required confidence for either class (see Fig SI2). This strategy mirrors the thresholding criteria used in AM [2].

### HbA Mutational Map

The final objective of this work was to construct a comprehensive mutational map of HbA based on predictions from the Combined Model (CM). This map is represented as an N × L matrix, where N = 20 corresponds to the number of possible SAS per residue, and L = 287 represents the total number of residues in HbA (considering one alpha and one beta chain). Thus, each value in the matrix represents a specific HbA variant, totaling 5,740 hypothetical variants. Cells are colored according to the predicted clinical significance: red for P, green for B, yellow for VUS, and gray for either empty cells or self-mutations. This visualization facilitates the identification of predictive patterns that support manual refinement of the model’s predictions (see Fig 7 and Fig 8). Rows, representing alternative amino acids, are ordered by chemical classification: aromatic, hydrophilic, hydrophobic, positively charged, or negatively charged. Columns, corresponding to reference amino acids, are ordered by structural classification (active site, core, surface, interface, and switch) and by preserving the relative residue order derived from the sequence alignment between the alpha and beta chains. In nearly half of the aligned residues between alpha and beta chains, structural classifications do not match. However, in most of these cases, one residue is assigned to the “switch” category. Given their sensitivity to conformational changes, switch residues are reassigned to match the structural category of their aligned counterpart, allowing for more consistent column ordering by structural and chemical criteria. Despite this adjustment, 15 aligned residue pairs remained incompatible, and 9 residues aligned to gaps in the MSA. In total, 39 residues (13.5% of all positions) were incompatible with simultaneous structural and chemical ordering. To prevent these from obscuring predictive patterns, they are grouped into a separate matrix section labeled “Incompatible”, preserving clarity in the main matrix (see Fig 8B).

This analysis revealed biologically meaningful patterns, allowing refinement of the CM predictions. The updated and corrected predictions are referred to as CM-PM (Combined Model - Post-Modelling).

A brief and illustrated guideline of the methodology recently described is presented below in Fig 1.

**Figure 1.**
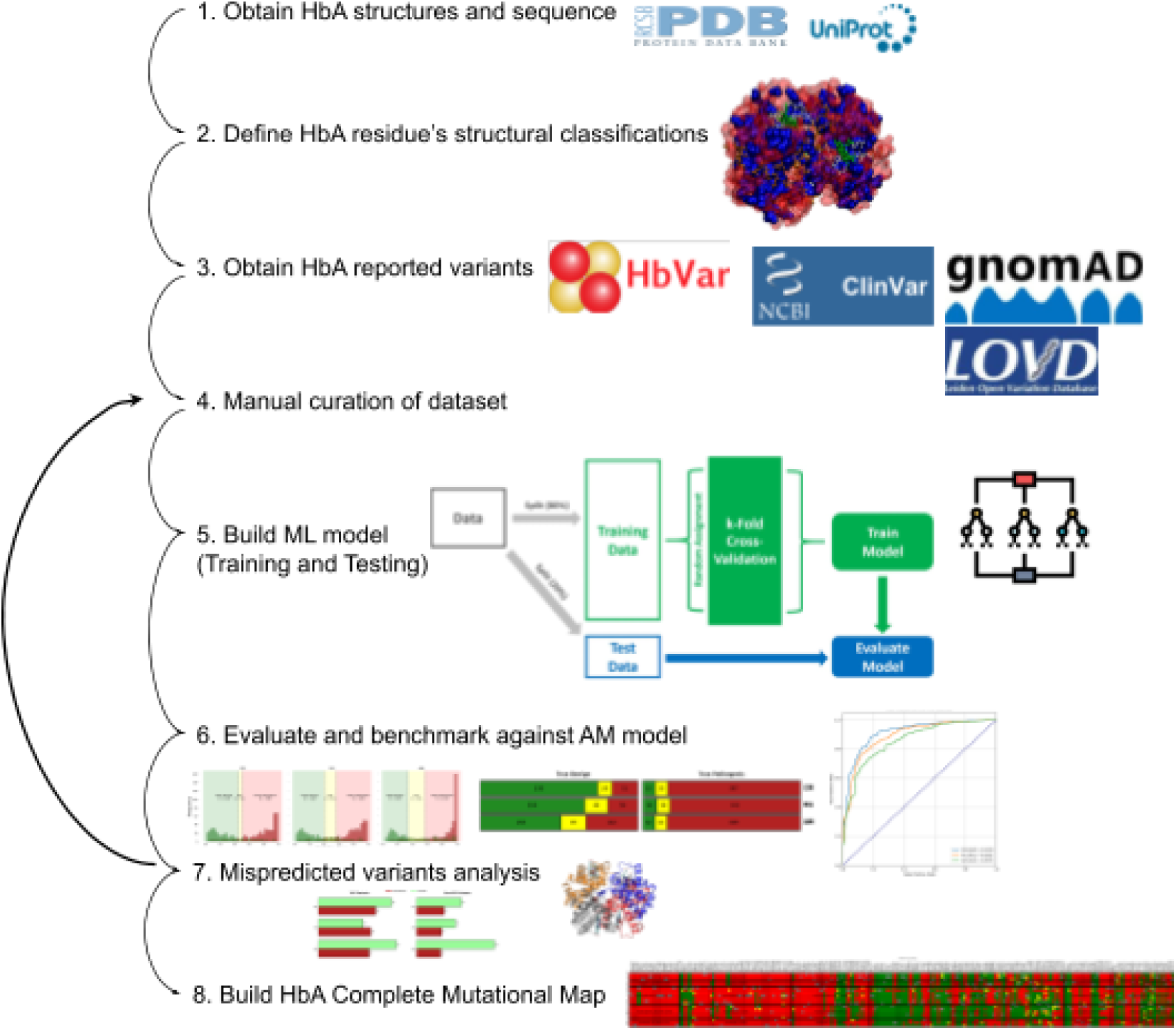
Methodology guideline illustration.

## Results

### Known Hb Variant Coverage

We began our analysis by assessing the coverage of available information (i.e. what is currently known) in relation to the total universe of potential HbA variants. Since our focus is at the protein level, we restricted the analysis to SAS. We considered two distinct sets of potential variants: (i) SAS resulting from a single base substitution (SBS) in the codon—i.e., only those amino acid changes that can arise from a single nucleotide mutation in the corresponding codon; and (ii) all theoretically possible SAS—i.e., any substitution of the wild-type residue with any of the other 19 standard amino acids, regardless of the underlying DNA mutation required. To assess available knowledge, we first analyzed whether the variants were classified as P, B, or VUS (see Methods). We further subclassified P variants by phenotype into the following categories: i) Unstable Hb, ii) Met-Hb, iii) altered oxygen affinity. The results are presented in Table 1.

**Table 1.**
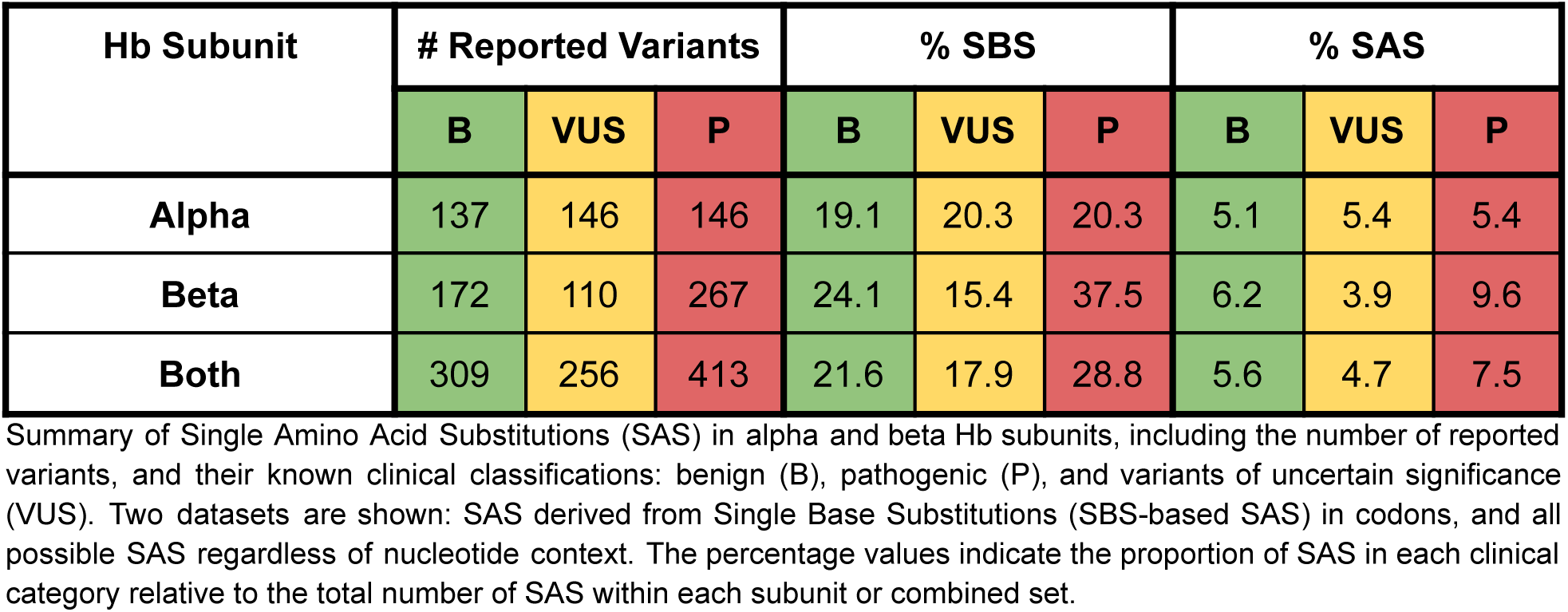
Hb variants and their known clinical classification.

Despite HbA being one of the most extensively studied proteins -and hemoglobinopathies among the earliest Mendelian diseases with a known molecular etiology-approximately half (ca. 695) of the possible SBS-derived SAS remains clinically unclassified. Moreover, when considering the full set of possible SAS, only ca. 15% have been clinically classified, leaving over 5000 variants as VUS. Clearly, even for this well-characterized protein and its associated phenotypes, there is still a need for reliable prediction tools for variant pathogenicity.

### Structural-Based Classification of Hb Variants

Hemoglobin structure-to-function relationships are possibly among the most thoroughly characterized in molecular biology. Therefore, we hypothesized that variant effects could be rationalized by analyzing their location and nature within the structural context of the protein, as well as the inherent Hb dynamics related to the T-to-R transition. In this context, a B variant would not disrupt the structure or dynamics of Hb, whereas P variants would likely produce measurable or significant alterations in any of them. We also reasoned that the type of effect should be mechanistically linked to the observed clinical phenotype. Thus, we hypothesized that: i) variants promoting heme autoxidation would lead to met-Hb, ii) variants destabilizing protein fold or core would cause unstable Hbs, iii) variants interfering with the R-to-T transition would alter oxygen affinity, and iv) variants reducing Hb solubility or promoting oligomerization would lead to sickle cell disease.

To support this structure-based interpretation of pathogenicity, we classified each residue in both the T and R states into five structural categories: i) Active Site (heme-contacting residues), ii) Core, iii) Surface, iv) Interface, and v) Switch residues (Fig 2). We then assessed the association between structural categories and both pathogenicity classification (P vs B) and individual phenotype classes. The results are presented in Fig 2.

**Figure 2.**
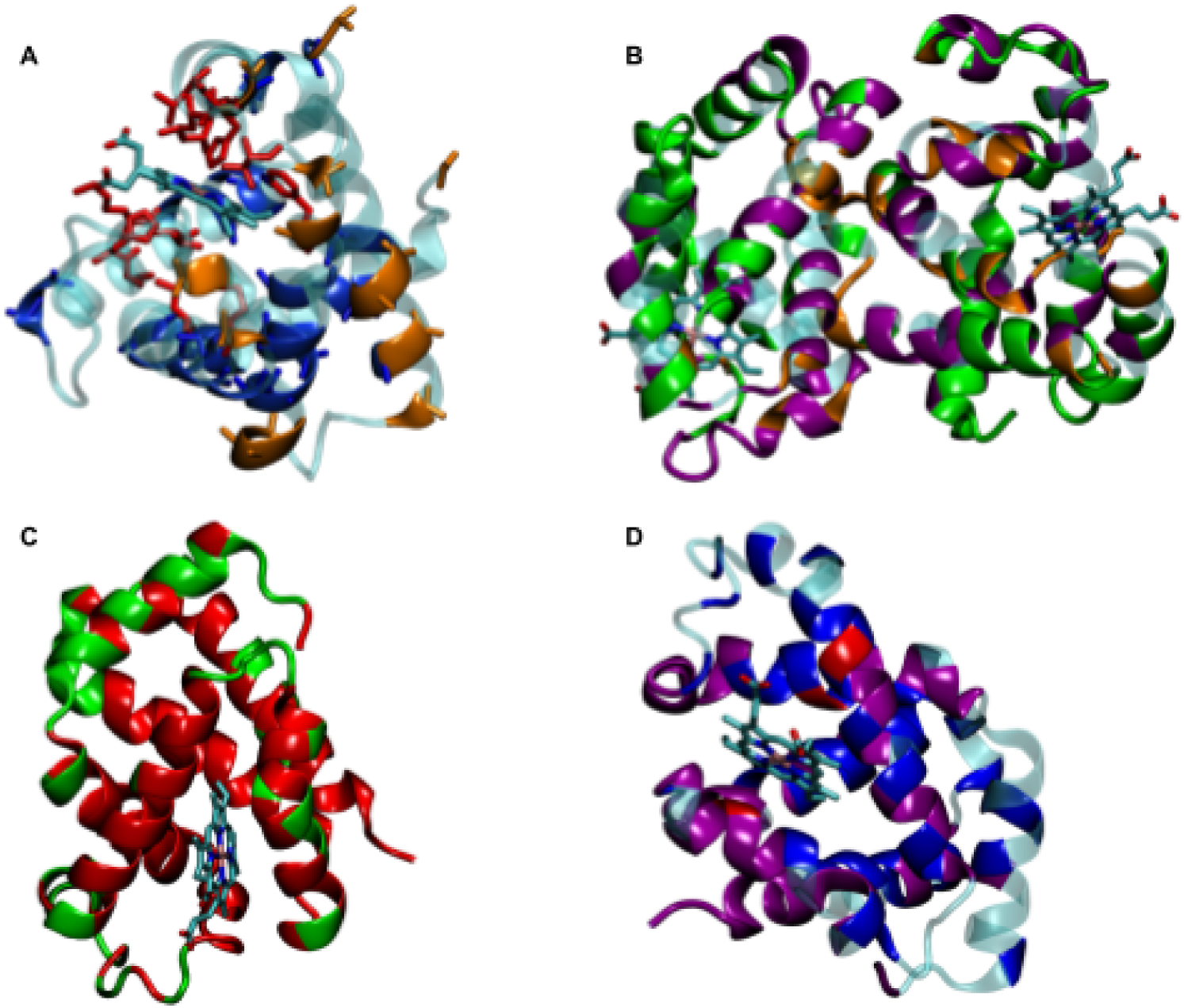
Structural classification of Hb residues on 1GZX structure **(A)** Mapping of Active Site (red), Core (blue), and Interface (orange) residues on the ∝-chain monomer. **(B)** Mapping of Interface (orange), Surface (green) and Switch (purple) residues on the ∝β dimer. **(C)** Mapping of residues predominantly associated with P (red) or B (green) variants on the β-chain monomer. **(D)** Mapping of β-chain residues associated with specific phenotypes: Unstable Hb (blue), altered O2 affinity (purple), met-Hb (red).

Fig 2 and Fig 3 show that the structural classification of Hb residues reveals clear trends in variant distribution and pathogenicity. Active site, core and interface residues show a strong enrichment for pathogenic variants. For instance, variants occurring in active site residues are approximately 10 times more likely to be pathogenic than benign, while core residues exhibit a 3-4-fold enrichment for pathogenicity. This trend is also visually evident in the structural mapping of the Hb subunits (compare Fig 2A and C). Switch residues, by contrast, appear to be more evenly distributed between benign and pathogenic variant classifications. Surface residues display the opposite bias, with benign variants outnumbering pathogenic ones by roughly 3 to 1. Looking at the structure, interface, switch, and surface residues (Fig 3B) are not easily distinguished.

**Figure 3.**
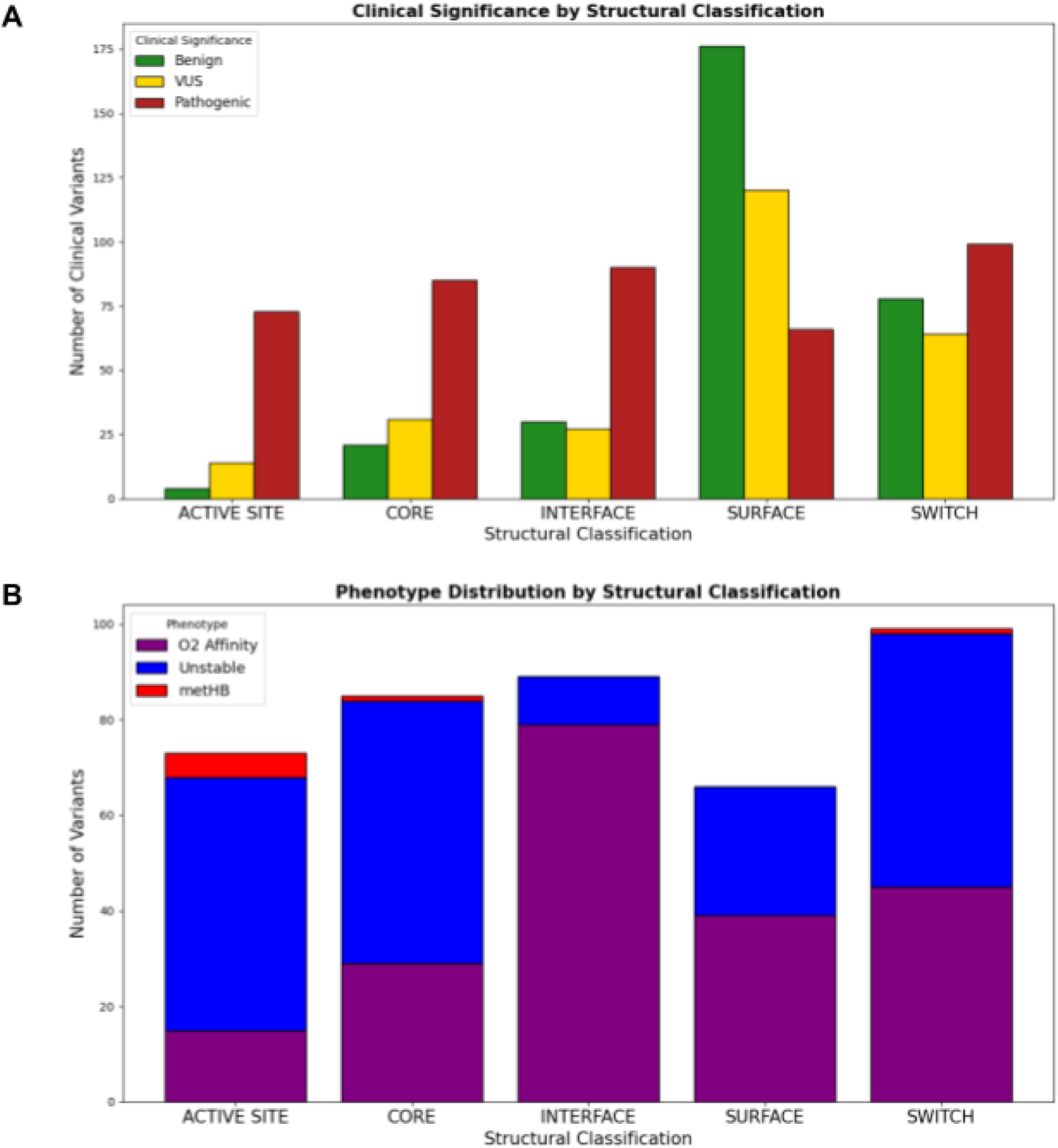
Distribution of curated variants across structural classifications. **(A)** Clinical significance categories. **(B)** Associated phenotypes.

Regarding phenotype-specific patterns, more subtle trends emerge. Unstable Hb variants are predominantly located in active site, core, and switch residues, with only a modest representation at the interface. In contrast, oxygen affinity–altering variants tend to localize at interface residues. MetHb-producing variants are exclusively found within the active site. The lower number of MetHb variants is likely due to their strict spatial restriction to the active site—a relatively small region of the protein—whereas other phenotypes can arise from variants spread across multiple structural regions. These structural associations provide useful insights into both variant classification and phenotype prediction.

### Analysis of Feature-pathogenicity Relationships

As already mentioned, for a SAS to result in a pathogenic variant and produce a specific phenotype, it must impact either the HbA structure or the dynamics of the T-to-R transition. To quantify these effects, we computed the following features: i) BLOSUM62 score, ii) FoldX ΔΔGfold, iii) ΔHydrophaty, iv) ΔMass, v) residue Conservation Score (rCS), vi) Chemical Mutation, vii) Topological Notation (i.e. The ∝-helix or loop in which the mutated residue lies) and viii) Structural Classification. We also included the AM score, which can be interpreted as a proxy for SAS effect, implicitly including structural context information.

We first analyzed each parameter distribution for both P and B variants to assess their predicting power. Properties based solely on residue substitution (such as BLOSUM62 score, ΔMass or ΔHydr) were not strongly informative and exhibited similar distributions for both classes (see Fig SI1). However, as expected -and with the exception of surface residues-P variants tend to show more negative BLOSUM62 scores (i.e. less conservative substitutions) whereas B variants cluster around neutral values.

FoldX ΔΔG emerged as a more effective pathogenicity predictor, specially when combined with

the rCS for the corresponding residue, as illustrated in the rCS vs FoldX ΔΔG plot (Fig 4A). This behavior is consistent with previous Massive mutagenesis Variant Experiments (MAVE) [4]. However, as shown in Fig 4A, there remains substantial overlap between P and B variants which justifies the need of more information to separate accordingly the P and B populations. The bar plots in Fig 4B, which displays the distribution of FoldX ΔΔG values across structural categories, indicate that although P variants generally have higher mean ΔΔG values, the distributions often overlap considerably. The most pronounced difference, as expected, is observed for core residues.

**Figure 4.**
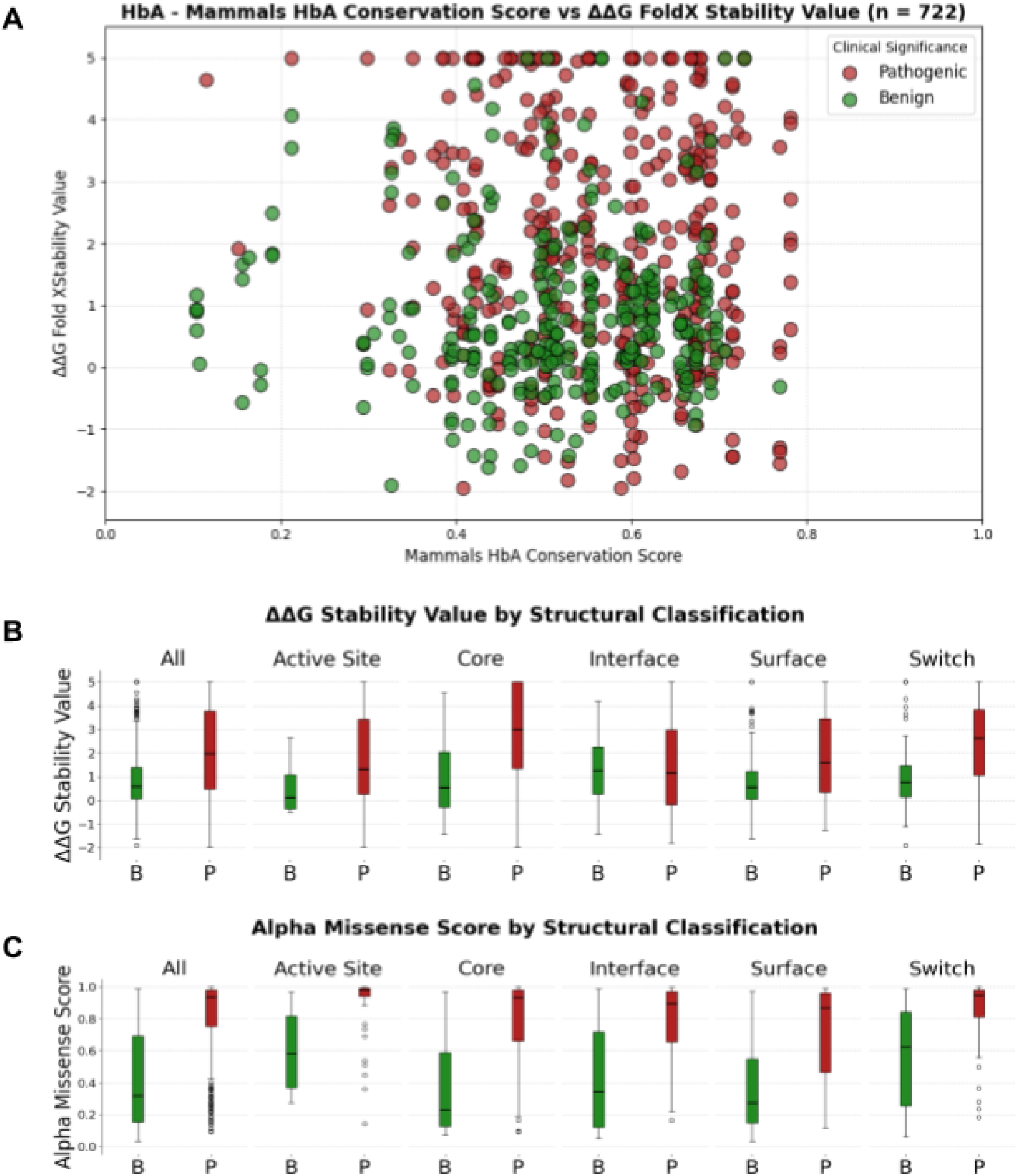
**(A)** Scatter plot of rCS vs FoldX ΔΔG for all P (red) and B (green) Hb variants. **(B)** Bar plot showing the distribution of FoldX ΔΔG values for P and B Hb variants. **(C)** Bar plot showing the distribution of AM scores for P and B variants.

The distribution of AM scores (Fig 4C) for HbA follows the expected overall trend, but also reveals several notable patterns. As expected, AM scores of P variants are clustered around high values, while B variants display a broader distribution skewed toward lower scores. Interestingly, the spread of AM score increases progressively across structural categories -from active site to core, interface and surface residues-mirroring the trend observed in Fig 3, where the likelihood of a variant being B increases along the same sequence. Interestingly, B variants at the active site tend to exhibit high AM scores. Together, these observations suggest that, although AM captures essential aspects of variant impact, it may have limitations in accurately identifying benign substitutions within functionally critical regions. Moreover, AM scores are generally skewed toward higher values, indicating a systematic tendency to overclassify variants as pathogenic.

Consistent with the observed results, no simple classification scheme based on a single feature -even when combined with structural classification-achieves reasonable accuracy in predicting variant effects. Therefore, in the following section, we develop and evaluate several ML models that integrate all available predictive features.

### Pathogenicity Prediction Model and Analysis

To build a pathogenicity prediction model, we employed ensemble-based methods (i.e. Random Forest-based Gboost) using all available variant-or residue-level features as predictors. Two models were developed: one excluding the AlphaMissense (AM) score (referred to as M1), and another including it (Combined Model, CM), in order to benchmark the contribution of AM as an external predictor. Analysis of ROC and Precision curves (Fig SI3) reveals that our M1 model already outperforms the standalone AM model (ROC AUC of 0.891 vs 0.857). Furthermore, incorporating the AM score as a feature allows a modest yet measurable improvement in performance (ROC AUC of 0.914).

Analysis of score distributions for known P and B variants (Fig SI4) reveals that the main difference between AM and our model lies in how scores are distributed, particularly among benign variants. For P variants, AM assigns predominantly high scores, while our model produces a similar pattern, though with generally lower scores -resulting in more variants falling near the classification threshold. In the case of benign variants, the difference is more pronounced: AM scores are slightly clustered toward lower values, whereas our model yields a rather flat Gaussian distribution slightly skewed toward lower scores.

The resulting predictions for each model are presented in Fig 5A as confusion matrices. All models perform similarly in identifying true pathogenic variants, with comparable true positive and false negative rates. However, the AM model exhibits a noticeably higher number of false positives and a substantially lower number of true negatives compared to both M1 and CM, suggesting lower overall specificity. In contrast, AM classifies fewer variants as VUS than M1, indicating a more decisive - though potentially overconfident - classification strategy. The CM model, which integrates the AM score as an additional feature, achieves the best overall performance. It preserves the sensitivity observed in the other models while improving specificity and reducing the number of variants classified as VUS -even fewer than AM-indicating a more confident and accurate classification. Although the improvement over M1 is not dramatic, the integration of AM clearly results in a more balanced and robust predictor.

**Figure 5.**
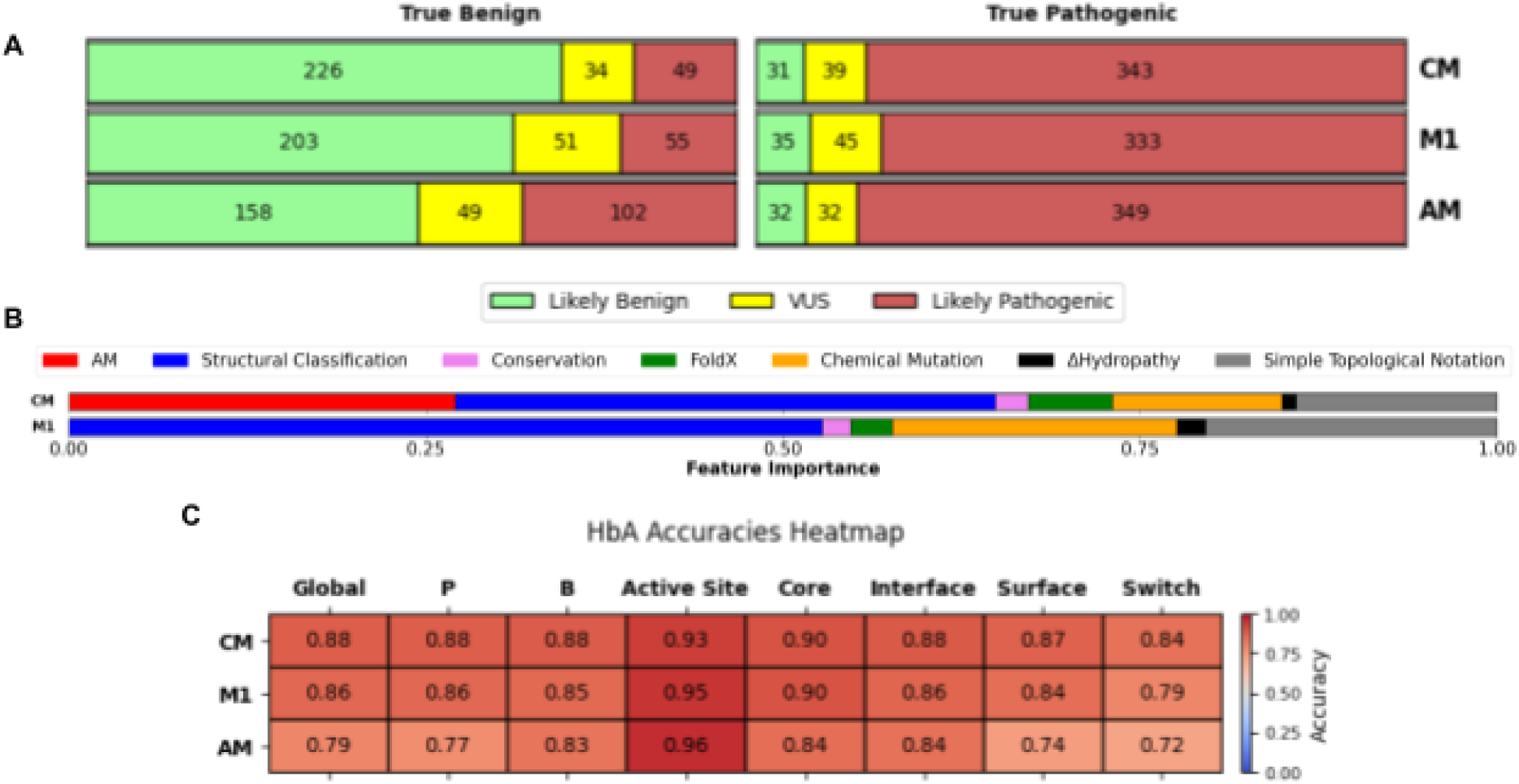
**(A)** Confusion matrix for the CM, M1 and AM models. **(B)** CM and M1 models features importances**. (C)** Models’ accuracies heatmap

Analysis of feature importance in our model (M1) reveals that structural classification is the most influential predictor, followed by topological notation -another structure-based property- and the Chemical Mutation features. This trend is maintained in the CM model, which additionally shows a significant contribution from the AM score (see Fig 5B). To further investigate this, we assessed the prediction accuracy of both AM and M1 across the different structural classification groups. As shown in Fig 5C, both methods exhibit a consistent performance trend across structural categories, while also revealing marked, category-dependent differences in accuracy. As expected, the active site shows the highest accuracy (>0.9), followed by core and interface residues, and lower values observed for surface and switch residues. The concordance between AM and M1 in accuracy profiles suggests that prediction difficulty is tied to the structural characteristics of each category. The switch region emerges as the most challenging for all models, likely due to its intrinsic structural flexibility and dynamic nature. While AM and M1 individually show reduced accuracy in this category, the CM model outperforms both, achieving superior performance. This suggests that integrating structural features with AM scores provides complementary insights that improve predictive accuracy in structurally ambiguous or heterogeneous contexts.

Another relevant observation is that, as previously discussed, the AM model tends to overpredict pathogenicity, assigning a disproportionately large number of variants to the pathogenic class. This bias is reflected in a markedly lower local accuracy for pathogenic variants compared to the M1 and CM models. In contrast, both structure-based models display a more balanced classification, effectively distinguishing between benign and pathogenic variants. This enhanced discrimination results in greater precision for pathogenic predictions and a reduction in false positives, reinforcing the benefit of incorporating structural features into the classification framework.

### Mispredicted Variants Analysis

Regarding misclassified variants, two main types of errors can be distinguished: (i) VUS misclassifications, in which variants known to be B or P are incorrectly classified as VUS, and (ii) non-VUS misclassifications, where the predicted classification is entirely incorrect (e.g., a benign variant predicted as pathogenic, or vice versa). Fig 6 summarizes the number of each misclassification type for each model.

**Figure 6.**
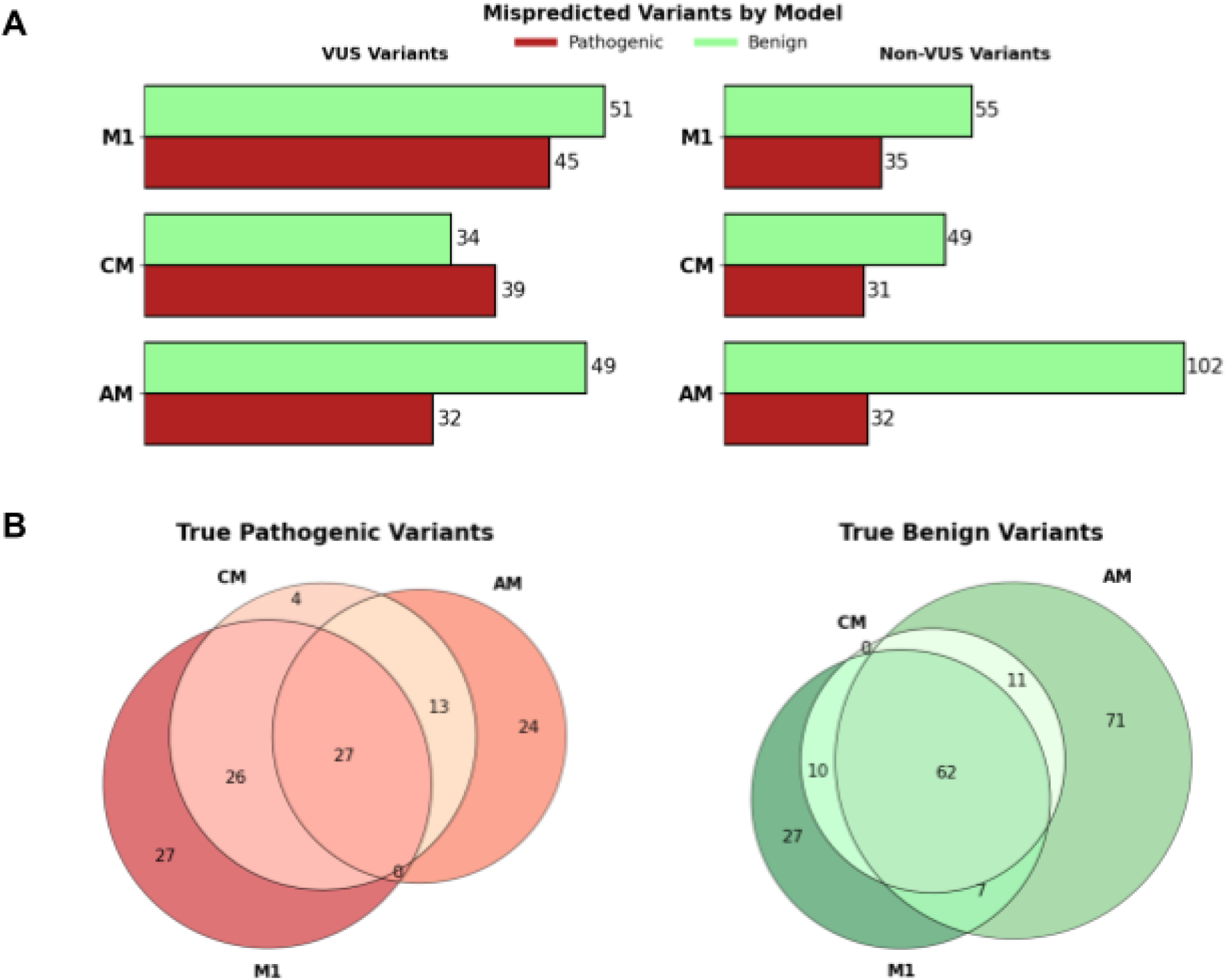
**(A)** Bar plots showing the number of mispredicted variants for each model for true P (red) and true B (green) variants. The left panel depicts VUS misclassifications while the right panel shows non-VUS misclassifications. **(B)** Venn Diagram illustrates the overlap of both VUS and non-VUS misclassifications among the CM, M1 and AM models for P (red, left) and B variants (green, right).

As shown in Fig 6A, although the AM model yields fewer VUS misclassifications compared to the M1 and CM, it exhibits a higher number of completely non-VUS misclassified variants. Notably, for non-VUS misclassifications, all methods tend to mispredict B variants more frequently than P variants. In contrast, for VUS misclassifications, while the AM model shows only a slight bias towards misclassifying B variants, the M1 and CM models more frequently fail to correctly classify P variants. However, the differences between misclassified B and P variants are small, and the lack of a clear trend suggests that no straightforward modification -such as threshold tuning-would lead to a significant improvement.

Fig 6B displays Venn diagrams illustrating the overlap of misclassifications (both VUS and non-VUS) for B and P variants separately. Notably, M1 and AM exhibit greater coincidence in misclassifying B variants compared to P variants (∼37% vs ∼22% out of all misclassified variants between both models on each class respectively). This moderate overlap suggests potential combination, whose positive effect is evidenced by the increased accuracy of the combined model (see Fig 5C). Moreover, the Venn diagrams show how the CM nicely blends AM and M1 but correctly covering more variants. These results suggest that further analysis could allow a better combination of both prediction methods to improve overall performance.

We now turn our attention to specific mispredicted examples to illustrate where the methods tend to fail, and to highlight ambiguities in clinical classification that will be further addressed in the next section. The first example is a false negative -predicted as benign but clinically classified as pathogenic-: Hb J-Cairo, a β-chain K>Q variant in the E8 position. The associated case report describes a mild phenotype, including moderately reduced cooperativity, slight microcytic anemia -possibly due to increased autoxidation-but no evidence of protein instability. Although clinically classified as pathogenic, the phenotypic impact is relatively modest. Both the AM and CM models incorrectly classify it as benign. Interestingly, the affected residue lies near the distal HisE7 and, although annotated as a surface residue, its structural coordinates fall just 0.1 units above the active site threshold -leading CM to classify it as pathogenic-. Its inclusion as an active site residue could be justified by the observed MetHb-like phenotype. Furthermore, the chemical nature of the substitution -changing a positively charged residue to a polar, uncharged one-has proven challenging for prediction, as approximately 50% of such mutations are mispredicted. The affected residue also holds a moderate rCS (0.617), which is not a strong indication of critical functional importance.

A second pathogenic variant misclassified by both models is Hb Calais, p.Ala77Pro substitution in chain β. This variant exhibits reduced oxygen affinity and elevated Met-Hb formation. Although the position is located in the key EF turn, the alanine residue is not highly conserved, and its substitution to proline does not induce significant destabilization (1.08 kcal/mol). Both methods fail to predict its pathogenicity, yielding scores of 0.571 and 0.537 for M1 and AM, respectively. Interestingly, although the MSA indicates low conservation of the alanine residue, proline is never observed at this position.

As a false positive case -a variant predicted as pathogenic by both models but classified as benign due to the absence of any clinical or hematological features-we analyzed Hb Kent, a WXXC substitution in the β-chain. Importantly, no biochemical evidence of hemoglobin instability was observed.. Analysis of the variant features reveals a high destabilizing effect (ΔΔG of ∼5.62 kcal/mol) and location within the C-helix and switch region -areas where most variants are pathogenic, though their mechanistic interpretation remains challenging. Visual inspection of the tetramer structure indicates that the Trp residue does not participate in any specific interactions, despite transitioning from the core (T-state) to the interface (R-state). In this case, the significant residue substitution and its structural location likely contribute to the pathogenic verdict; however, the lack of specific functional interactions seems to result in no observable effect.

A more subtle case of a variant misclassified as pathogenic is Hb Geisinger, an ∝-chain p.Phe37Leu mutation. The residue is structurally classified as switch, transitioning from interface (R) to core (T) region, with a ΔΔG of 2.73, suggesting a potentially destabilizing role that may explain its pathogenic prediction. Clinically, this variant is reported as benign; however, the patient exhibits symptoms of anemia, which are attributed to a deletion in the HBA1 gene by the authors. Interestingly, the variant is absent from gnomAD.

### Proposed Variant Classification Revisions

In the course of the above misprediction analysis we identified several cases for which there is strong evidence suggesting a potentially incorrect clinical classification, as currently listed in HbVar and/or ClinVar. Examples are provided below:

The p.Ala11Thr substitution in the β-chain, known as Hb Belleville, involves the replacement of a hydrophobic alanine with a hydrophilic threonine within an ∝-helix. Both the CM and AM models predict this variant to be benign. Despite its current classification in HbVar as pathogenic -due to its association with anemia-a closer examination supports a benign interpretation. The AM score suggests a minor effect (0.0768), consistent with FoldX showing no significant destabilization in folding energy (0.582 kcal/mol). The variant resides in the switch region, where both B and P variants are found, and visual inspection of the structure reveals no relevant interactions involving the affected residue. Therefore, the model prediction is consistent and well supported. Interestingly, the case report linked to this variant attributes the observed anemia to chronic diabetes, rather than to the Hb mutation itself. Biochemical stability assays yielded a negative isopropanol test and a mildly positive heat test. Moreover, in ClinVar the clinical significance of this variant is labeled as “Conflicting Interpretations of Pathogenicity”, with one submission classifying it as Likely Benign and another one as “Uncertain significance”. Additionally, gnomAD reports the presence of four alleles (allele frequency of 2.5 *x* 10^−6^). Taken together, these data together support a reclassification of this variant as benign.

Another example of a variant originally labeled as pathogenic but reclassified as benign based on model predictions is Hb Harbin, a K17M substitution in the ∝-chain (HBA1/2). Listed in HbVar as associated with anemia, the variant displays an increased O2 affinity and mild instability. The affected residue lies on the protein surface, where the mutation replaces a positively charged, hydrophilic lysine with a hydrophobic methionine. Although such substitutions can potentially reduce solubility or interfere with interactions at the aqueous interface, this is not necessarily the case. In this particular context, K17 does not engage in any critical interactions or structural networks, and neighboring residues do not appear to depend on its polarity for structural stability or function. Both the FoldX ΔΔG value (−0.23 kcal/mol) and the rCS (0.51) support a benign interpretation, indicating minimal structural impact and evolutionary tolerance. This variant is not reported in gnomAD. Closer examination of the original publication describing Hb Harbin [3], shows that although originally, the variant tested positive for both increased oxygen affinity and reduced stability, it was also identified in other Chinese families without any associated clinical symptoms. Based on these observations, the authors concluded that the clinical abnormalities observed in their patient -such as anemia-were likely attributable to other unrelated causes rather than the HbA variant itself. This revised interpretation is further supported by the pathogenicity score from our CM model (0.12), which aligns with a benign classification and suggests that the previous pathogenic label assigned to Hb Harbin may have been inaccurate. Consistently, Hb Harbin is also predicted as benign by the AM model, with a score of 0.29. Moreover, all other variants reported in HbVar at position K17 have been associated with no clinical phenotype, including Hb Boa Esperança (p.Lys17Thr), Hb I (p.Lys17Glu), and Hb Beijing (p.Lys17Asn). This pattern suggests that K17 is a permissive site, capable of accommodating amino acid substitutions without leading to pathological consequences.

Two related variants displaying opposite classification behaviour are p.Arg142Pro (Hb Singapore) and p.Asp127Ala (Hb Verdun). Both are currently classified as Benign but were predicted by both models as pathogenic with high confidence. Structurally, both residues lie at the α/α interface, establishing a salt bridge between them. Mutation of either residue disrupts this interaction. Analysis of variant characteristics show a moderate conservation (rCS between 0.5-0.7) and slight destabilization (ΔΔG = 2.15 kcal/mol), suggesting a potential structural impact. Clinical evidence, however, is less conclusive. In the case of Hb Singapore, although no abnormal laboratory findings were reported, the patient presented with mild anemia that responded to iron therapy. For Hb Verdun, no detailed clinical presentation is provided, and its classification as Benign is based solely on normal hematological parameters in a heterozygous individual, in whom the mutant Hb represented only 17% of the total Hb. Most importantly, another variant at the same position, p.Asp127Val (Hb Fukutomi) has been reported as pathogenic due to altered O2 affinity. Similarly, in the HBA2 gene, a variant at the same residue—p.Asp127Asn (Hb Tarrant)—has also been associated with pathogenicity for the same reason. As in previous cases, the overall evidence -including predictive model scores and careful review of the clinical context-suggests that both variants should be reclassified as pathogenic.

### A Complete Mutational Map of HbA

To draw a comprehensive view of the HbA mutational landscape that could provide a final refinement of its predictive capacity, we built and analyzed different SAS matrices. In these matrices, the x-axis represents the residue number and the y-axis the 20 possible SAS. Each cell can be coloured in green or red for B or P variants respectively and also yellow for VUS variants. Additionally, each cell may be splitted in two triangles if the matrix is used to contrast the reported clinical significance of the variants against its prediction by the CM model, in this case the lower triangle represents the predicted output while the upper triangle represents the real reported pathogenicity. Given the relevance of residue structural classification, residues from both ∝ and β chains were reordered according to their corresponding structural categories. Structurally equivalent ∝ and β residues were placed adjacent to one another, based on a pairwise sequence alignment between the two chains. Thus, the columns are not only ordered by structural classification, but also aligned across chains according to their sequence correspondence. However, there are a few exceptions where residues from one chain do not have a matching counterpart in the other, or where they do match but their structural classifications do not coincide. In such cases, if either residue is classified as part of the Switch region—a structurally dynamic zone—this mismatch is tolerated and they are still considered structurally equivalent. Despite this criterion, a total of 39 residues (∼13.5%) could not be confidently aligned and classified. These residues were grouped into a marginal structural category named ‘Incompatible’, which does not follow a consistent logical ordering and is therefore excluded from the pattern analysis in the mutational map. The column labels are the residue’s chain followed by its one letter amino acid code and its position on the protein sequence.

Fig 7 displays the SAS matrix for clinically annotated variants alongside their corresponding predictions by the CM model on the switch region. The matrix highlights the limited current knowledge regarding the potential universe of SAS, as well as emerging trends in variant type (B vs P) and model prediction accuracy across different structural regions. Most active site variants are pathogenic and correctly predicted, whereas surface residues tend to harbor more benign variants, for which prediction is less accurate (see Fig 8 and SI).

**Figure 7.**
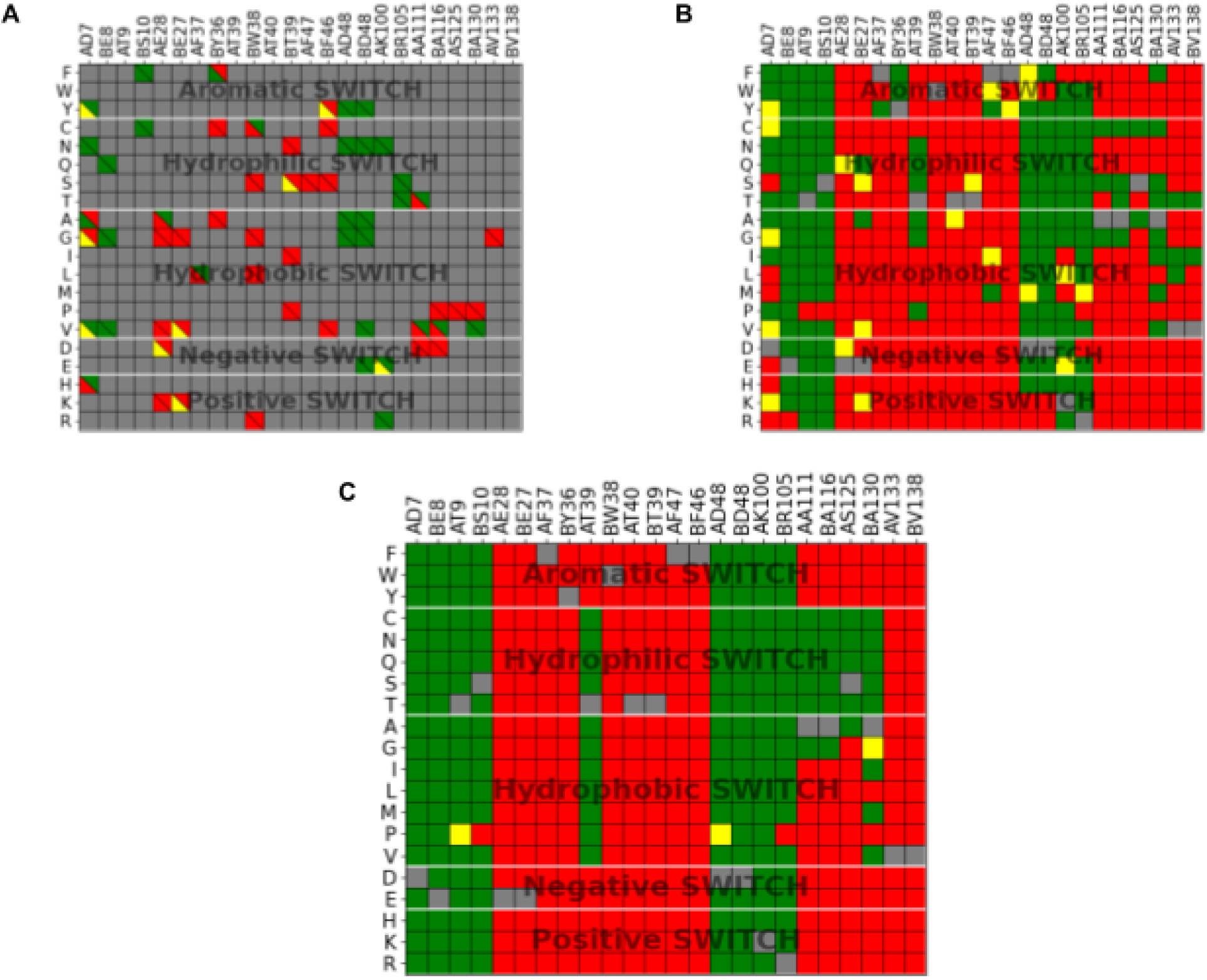
Mutational maps for the switch region. **(A)** Comparison between the clinical significance of reported variants (upper triangles) and CM predictions (lower triangles). Green indicates benign, red pathogenic, yellow VUS, and grey denotes unreported variants, excluded variants, or synonymous mutations. **(B)** Full mutational map showing all CM predictions for the switch region. **(C)** Final mutational map of CM-PM predictions, incorporating manually defined correction patterns based on observations from panel B.

**Figure 8.**
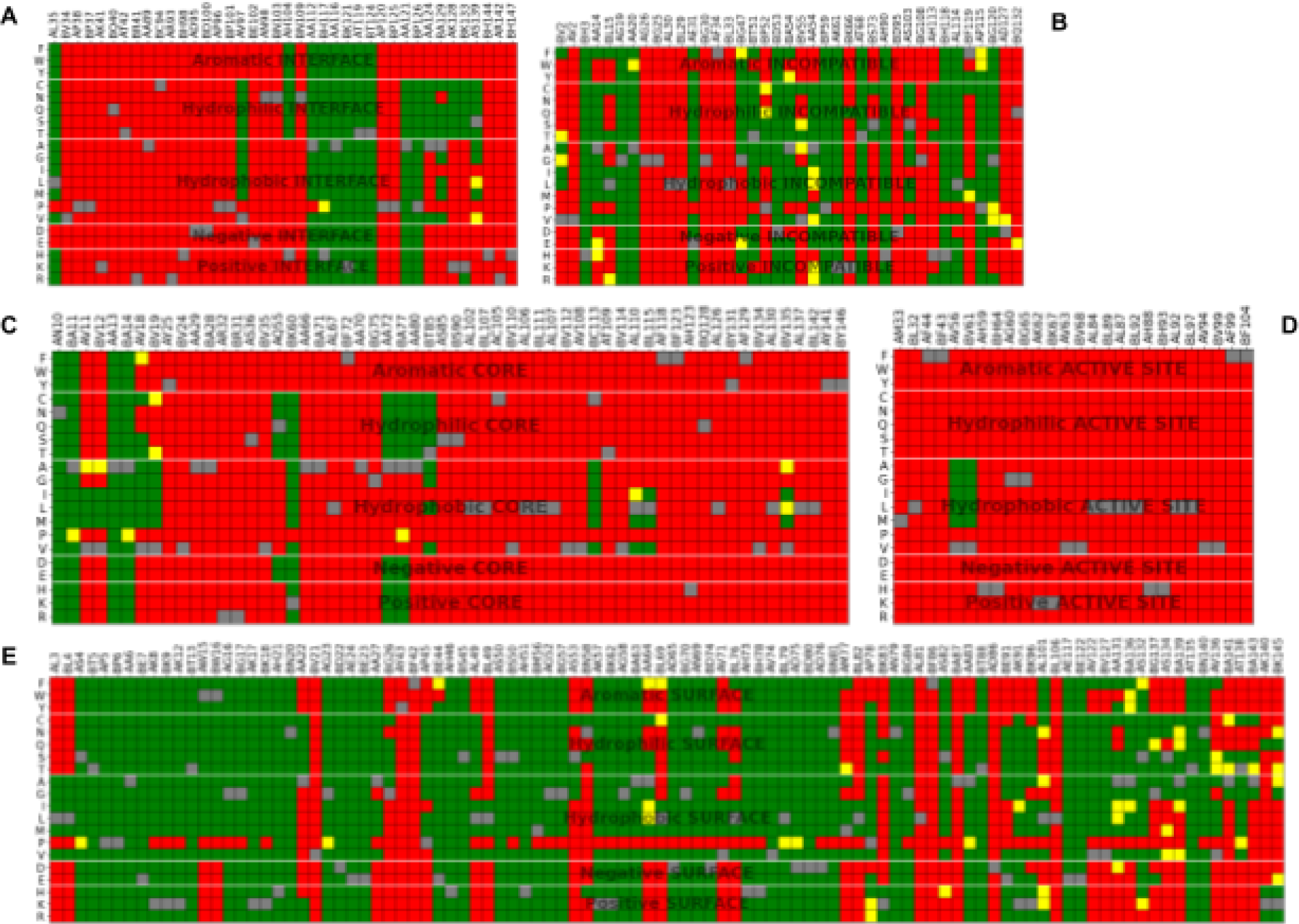
Final mutational maps of CM-PM predictions for the **(A)** Interface **(B)** Incompatible **(C)** Core **(D)** Active Site and **(E)** Surface regions. The incompatible region is not analyzed nor modified since its residues are not logically ordered by its columns, thus coinciding with its initial CM prediction’s mutational map.

Building the complete SAS matrix (i.e. the final mutational map of CM-PM predictions) based on CM predictions (Fig 7C and Fig 8) reveals even more notable outcomes. The matrix exhibits clear square-like patterns of consistent predictions, corresponding to groups of residues sharing the same structural classification, topology and mutation type, which collectively lead to uniform predictions. For many positions, the predictions are consistent across all possible SAS (i.e., columns displaying a single color), while within the same structural classification group, sub-regions with distinct prediction patterns are evident. Additionally, clusters of adjacent columns display rectangular zones where substitutions to chemically similar residues yield either benign or pathogenic predictions. Note that we deliberately excluded variants with proline substitutions due to their unpredictable and context-dependent structural effects.

We investigated whether the observed rectangular patterns in the SAS matrix could be used to improve the accuracy of B/P variant prediction. Assuming that these patterns reflect biologically meaningful consistency, we implemented a post hoc correction by enforcing uniform predictions within a given rectangular region whenever more than X% of the SAS shared the same predicted outcome and aligned with established clinical classifications. As an illustrative example, Fig 7 displays the matrix section corresponding to adjacent residues AD48, BD48, AK100 and BR105 from the switch region. In the CM matrix, several substitutions are correctly predicted as benign; however, a number of pathogenic predictions appear to be erroneous, and many variants are classified as VUS. In contrast, the PM model consistently classifies the entire set of substitutions as benign—except for those involving proline. Structural inspection of these residues confirms that they are all quite far from the heme iron and oriented towards the protein surface, supporting the benign classification.

The primary impact of the CM-PM model is a significant reduction in the number of variants classified as VUS (from 73, ∼10.1%, to 20, ∼2.8%), with a vast majority of these now correctly predicted (see Fig 9). More importantly, the number of true positives and negatives increases while the number of false positives and negatives decreases. This improvement is reflected in a modest overall increase in accuracy from 0.88 to 0.89, driven by gains across all structural classes, particularly within the switch region. The only exception is the active site, where a slight decrease from 0.93 to 0.91 in accuracy is observed (see Fig SI5).

**Figure 9.**
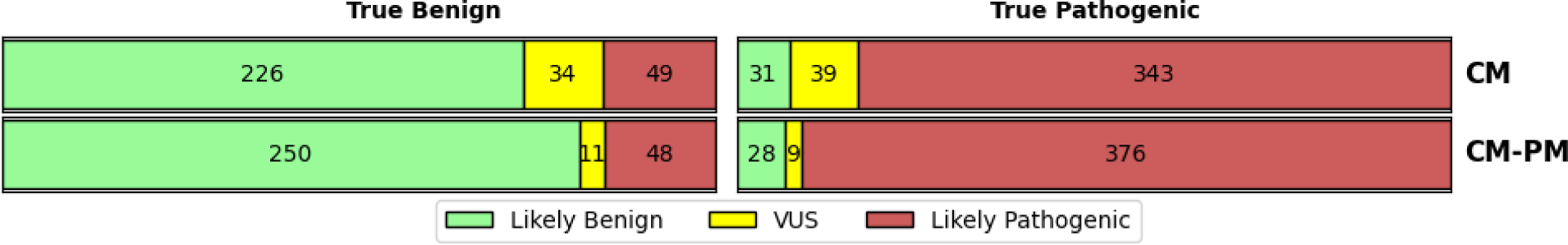
Confusion matrix of CM model before (CM) and after HbA mutation map refinement (CM-PM)

## Discussion

The central hypothesis of this study is that a comprehensive understanding of the interplay between protein sequence, structure/dynamics, and function can lead to clearer insights into the mechanisms driving pathogenicity, ultimately enabling the development of rational and interpretable models for variant classification. Our findings provide strong support for this premise. Early analyses revealed distinct trends linking structural context to clinical outcome, which laid the groundwork for the development of a predictive model capable of accurately distinguishing between pathogenic and benign variants in HbA. Notably, the model’s performance strongly relies on the structural classification of the mutated residue.

Our results demonstrate that knowledge-based structural features are powerful predictors of pathogenicity. In benchmarking experiments, our model achieved performance metrics comparable to -and in some cases exceeding-those of widely used general pathogenicity predictors, such as AlphaMissense. Moreover, we observed that combining our structure-informed model with AM led to synergistic improvements in classification accuracy. These findings suggest that complementing or fine-tuning data-driven models trained on broader genomic contexts can significantly improve both the accuracy and interpretability of pathogenicity predictions.

The provided analytical framework also demonstrates practical utility in addressing clinically ambiguous cases. We re-evaluated several variants of uncertain significance or with conflicting annotations and -based on their structural and functional contexts-proposed reclassifications supported by the model’s predictions. These results underscore the potential of mechanistically grounded tools to enhance the clinical interpretation of variants, and will help to improve the overall accuracy of clinical information available in public databases.

We additionally generated a comprehensive mutational landscape of HbA, predicting the effect of every possible SAS. This resource offers high-resolution insights into the structural and functional tolerance of HbA, which may aid in the interpretation of novel variants and potentially inform future experimental studies. To support transparency and reproducibility, both the trained model and the resulting mutational map have been made freely available to the research and clinical communities (See SI).

It is also important to remark that, while our approach shows considerable promise, it has certain limitations. The current model is specific to HbA, and its performance on other proteins may depend on the availability and quality of structural data, as well as on the depth of clinical knowledge regarding SAS. Furthermore, although we incorporated a broad range of features, some of them -such as the switch region classification-may only apply to other allosteric systems, while others, including post-translational modifications, may require protein-specific fine-tuning. Nonetheless, we expect that future work will expand this framework to additional clinically relevant proteins and explore the integration of additional data sources, such as evolutionary couplings, intraprotein residue networks and protein-protein interaction data.

We also aim to further enhance the interpretability of the model, enabling clinicians to better understand the specific structural or biochemical features underlying individual phenotype-specific predictions. In the present study, the limited number of variants associated with each phenotype precluded the development of phenotype-specific ML models. As a result, only general trends and associations could be derived (Fig 2 and Fig 3).

In summary, our findings validate the central hypothesis and underscore the value of integrating protein-level mechanistic insights into variant pathogenicity prediction. This structure-informed and interpretable framework holds potential for improving diagnostic accuracy and enhancing our understanding of the molecular mechanisms underlying disease-causing variants.

## Supporting information

Supplementary Image 1

Supplementary Image 2

Supplementary Image 3

Supplementary Image 4

Supplementary Image 5

HbA Variants Data

## Acknowledgements

The authors gratefully acknowledge the Facultad de Ciencias Exactas y Naturales, Universidad de Buenos Aires (FCEyN-UBA) for providing the facilities, resources, and academic environment that made this work possible. We are especially thankful for the financial support received through the Becas UBACyT Estímulo program, which enabled and sustained the development of this research.

## References

1. Valdar WS. Scoring residue conservation. Proteins. 2002 Aug 1;48(2):227-41. doi: 10.1002/prot.10146. PMID: 12112692.

2. Cheng J, Novati G, Pan J, Bycroft C, Žemgulytė A, Applebaum T, Pritzel A, Wong LH, Zielinski M, Sargeant T, Schneider RG, Senior AW, Jumper J, Hassabis D, Kohli P, Avsec Ž. Accurate proteome-wide missense variant effect prediction with AlphaMissense. Science. 2023 Sep 22;381(6664):eadg7492. doi: 10.1126/science.adg7492. Epub 2023 Sep 22. PMID: 37733863.

3. Zeng YT, Huang SZ, Qiu XK, Cheng GC, Ren ZR, Jin QC, Chen CY, Jiao CT, Tang ZG, Liu RH, et al. Hemoglobin Chongqing [alpha 2(NA2)Leu----Arg] and hemoglobin Harbin [alpha 16(A14)Lys Met] found in China. Hemoglobin. 1984;8(6):569-81. doi: 10.3109/03630268408991742. PMID: 6526652.

4. Høie, Magnus & Cagiada, Matteo & Frederiksen, Anders & Stein, Amelie & Lindorff-Larsen, Kresten. (2022). Predicting and interpreting large-scale mutagenesis data using analyses of protein stability and conservation. Cell Reports. 38. 110207. 10.1016/j.celrep.2021.110207.

5. Schymkowitz, Joost et al. “The FoldX web server: an online force field.” Nucleic acids research vol. 33,Web Server issue (2005): W382–8. doi:10.1093/nar/gki387

6. Hardison RC, Chui DH, Giardine B, Riemer C, Patrinos GP, Anagnou N, Miller W, Wajcman H. HbVar: A relational database of human hemoglobin variants and thalassemia mutations at the globin gene server. Hum Mutat. 2002 Mar;19(3):225–33. doi: 10.1002/humu. 10044. PMID: 11857738.

7. Landrum, Melissa J et al. “ClinVar: public archive of relationships among sequence variation and human phenotype.” Nucleic acids research vol. 42,Database issue (2014): D980–5. doi:10.1093/nar/gkt1113

8. Karczewski, Konrad J et al. “The mutational constraint spectrum quantified from variation in 141,456 humans.” Nature vol. 581,7809 (2020): 434–443. doi:10.1038/s41586-020-2308-7

9. Fokkema, Ivo F A C et al. “The LOVD3 platform: efficient genome-wide sharing of genetic variants.” European journal of human genetics : EJHG vol. 29,12 (2021): 1796–1803. doi:10.1038/s41431-021-00959-x

10. Bringas M, Petruk AA, Estrin DA, Capece L, Martí MA. Tertiary and quaternary structural basis of oxygen affinity in human hemoglobin as revealed by multiscale simulations. Sci Rep. 2017 Sep 7;7(1):10926. doi: 10.1038/s41598-017-11259-0. PMID: 28883619; PMCID: PMC5589765.

11. Schuster CD, Salvatore F, Moens L, Martí MA. Globin phylogeny, evolution and function, the newest update. Proteins. 2024 Jun;92(6):720–734. doi: 10.1002/prot.26659. Epub 2024 Jan 9. PMID: 38192262.

12. Jorge SE, Bringas M, Petruk AA, Arrar M, Marti MA, Skaf MS, Costa FF, Capece L, Sonati MF, Estrin D. Understanding the molecular basis of the high oxygen affinity variant human hemoglobin Coimbra. Arch Biochem Biophys. 2018 Jan 1;637:73–78. doi: 10.1016/j.abb.2017.11.010. Epub 2017 Dec 1. PMID:29199120.

13. Song S, Starunov V, Bailly X, Ruta C, Kerner P, Cornelissen AJM, Balavoine G. Globins in the marine annelid Platynereis dumerilii shed new light on hemoglobin evolution in bilaterians. BMC Evol Biol. 2020 Dec 29;20(1):165. doi: 10.1186/s12862-020-01714-4. PMID: 33371890; PMCID: PMC7771090.

14. Bustamante JP, Radusky L, Boechi L, Estrin DA, Ten Have A, Martí MA. Evolutionary and Functional Relationships in the Truncated Hemoglobin Family. PLoS Comput Biol. 2016 Jan 20;12(1):e1004701. doi: 10.1371/journal.pcbi.1004701. PMID: 26788940; PMCID: PMC4720485.

15. Wisecaver JH, Alexander WG, King SB, Hittinger CT, Rokas A. Dynamic Evolution of Nitric Oxide Detoxifying Flavohemoglobins, a Family of Single-Protein Metabolic Modules in Bacteria and Eukaryotes. Mol Biol Evol. 2016 Aug;33(8):1979–87. doi: 10.1093/molbev/msw073. Epub 2016 Apr 12. PMID: 27189567.

16. Embury, Stephen H. Sickle Cell Disease : Basic Principles and Clinical Practice. New York: Raven Press, 1994. Print.

17. UniProt Consortium, The. “UniProt: the universal protein knowledgebase.” Nucleic acids research vol. 46,5 (2018): 2699. doi:10.1093/nar/gky092

18. Crystallography: Protein Data Bank. Nature New Biology 233, 223 (1971). 10.1038/newbio233223b0

19. Cock, Peter J A et al. “Biopython: freely available Python tools for computational molecular biology and bioinformatics.” Bioinformatics (Oxford, England) vol. 25,11 (2009): 1422–3. doi:10.1093/bioinformatics/btp163

20. Katoh, Kazutaka et al. “MAFFT: a novel method for rapid multiple sequence alignment based on fast Fourier transform.” Nucleic acids research vol. 30,14 (2002): 3059–66. doi:10.1093/nar/gkf436

21. Henikoff, S, and J G Henikoff. “Amino acid substitution matrices from protein blocks.” Proceedings of the National Academy of Sciences of the United States of America vol. 89,22 (1992): 10915–9. doi:10.1073/pnas.89.22.10915

22. Stone, M. “Cross-Validatory Choice and Assessment of Statistical Predictions.” Journal of the Royal Statistical Society. Series B (Methodological), vol. 36, no. 2, 1974, pp. 111–47. JSTOR, http://www.jstor.org/stable/2984809. Accessed 19 June 2025.

23. Breiman, Leo. “Random forests.” Machine learning 45 (2001): 5–32.

24. Cortes, C. and Vapnik, V. (1995) Support-Vector Networks. Machine Learning, 20, 273–297. 10.1007/BF00994018

25. Chen, T., and Guestrin, C. (2016). XGBoost: A scalable tree boosting system. Proceedings of the 22nd ACM SIGKDD International Conference on Knowledge Discovery and Data Mining. 10.1145/2939672.2939785

