## Supplementary figures and images for "A structure and function-based complete mutational map of Human Hemoglobin using AI"

### Supplementary Image 1

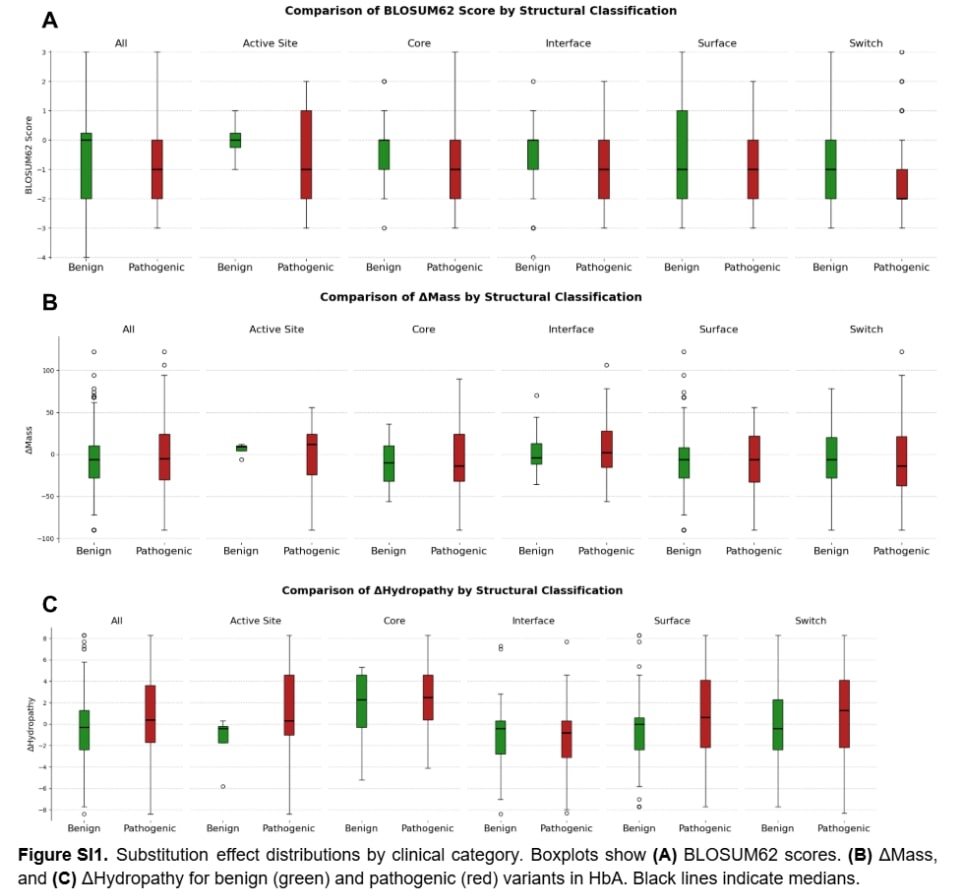

### Supplementary Image 2

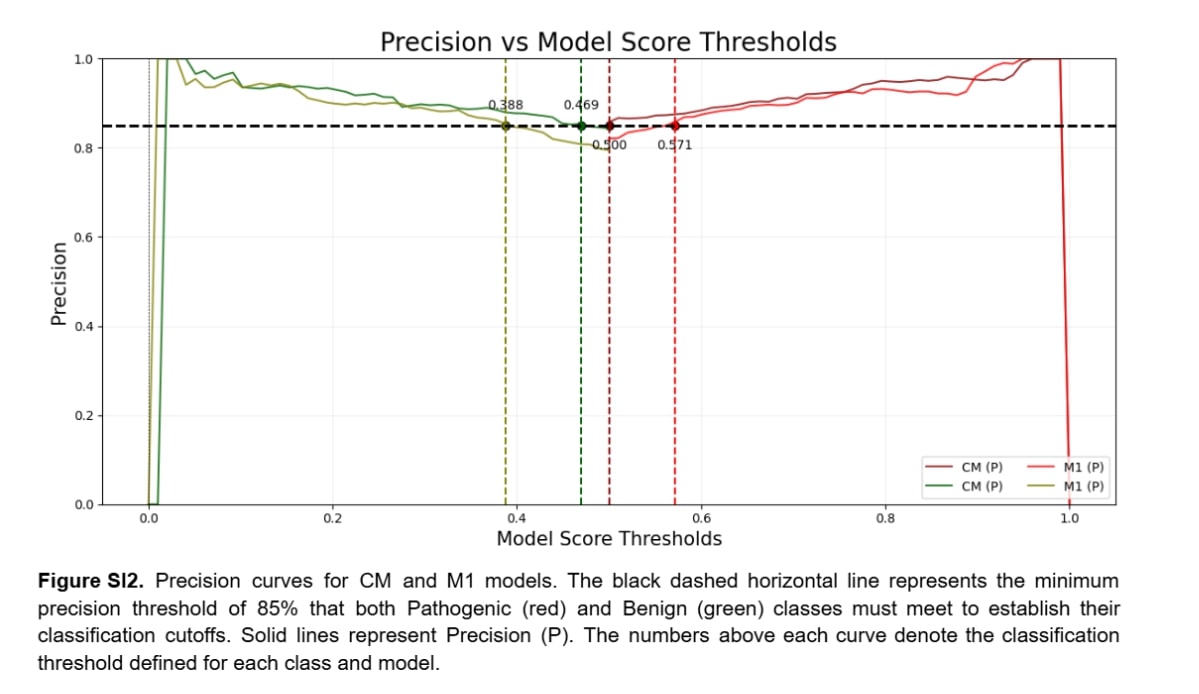

### Supplementary Image 3

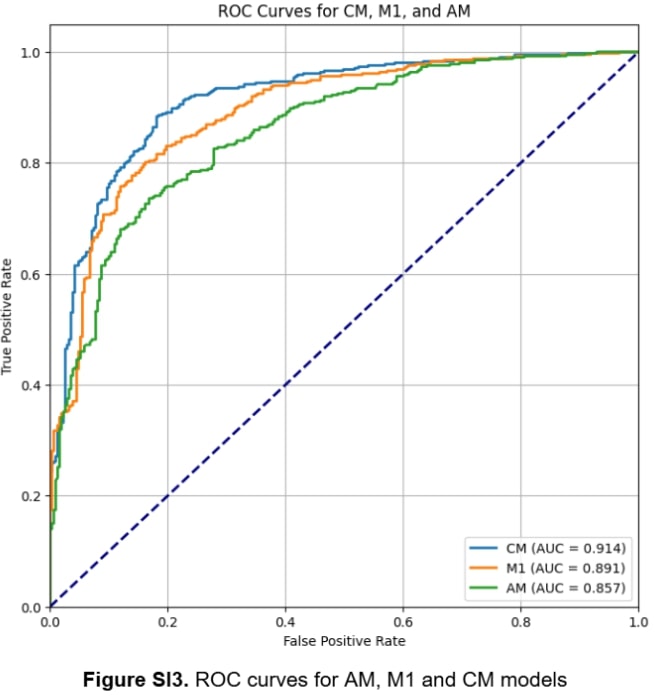

### Supplementary Image 4

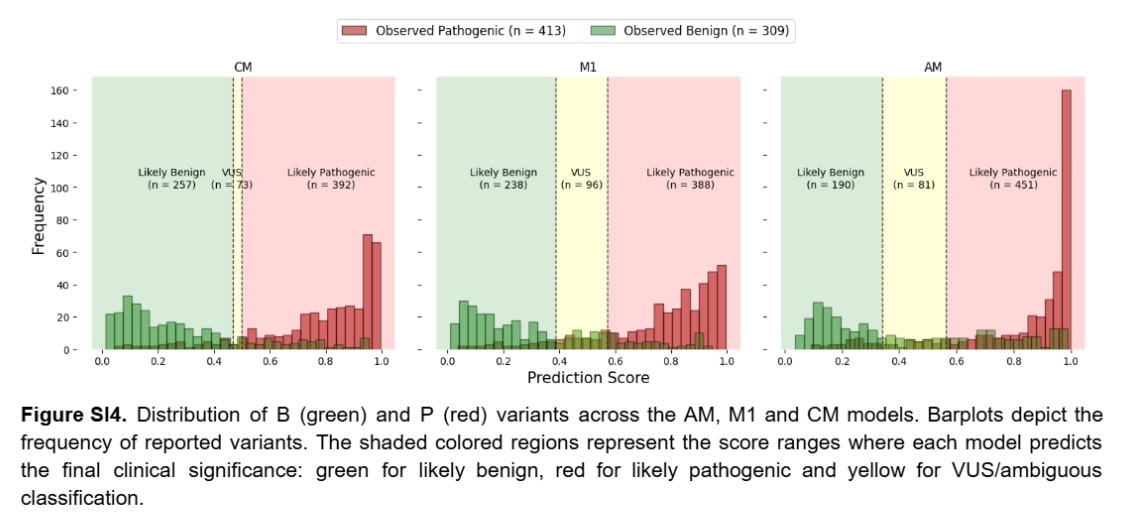

### Supplementary Image 5

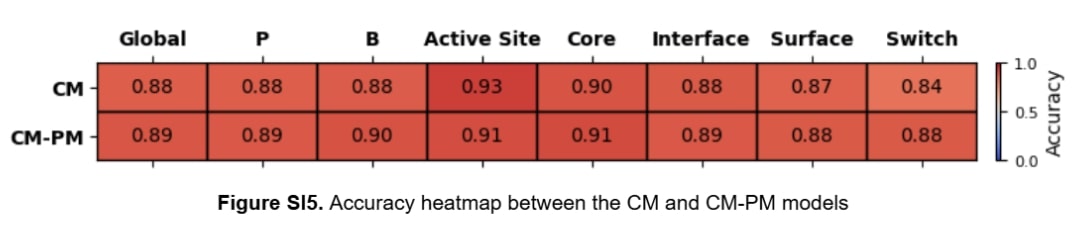

### Supplementary Table 1

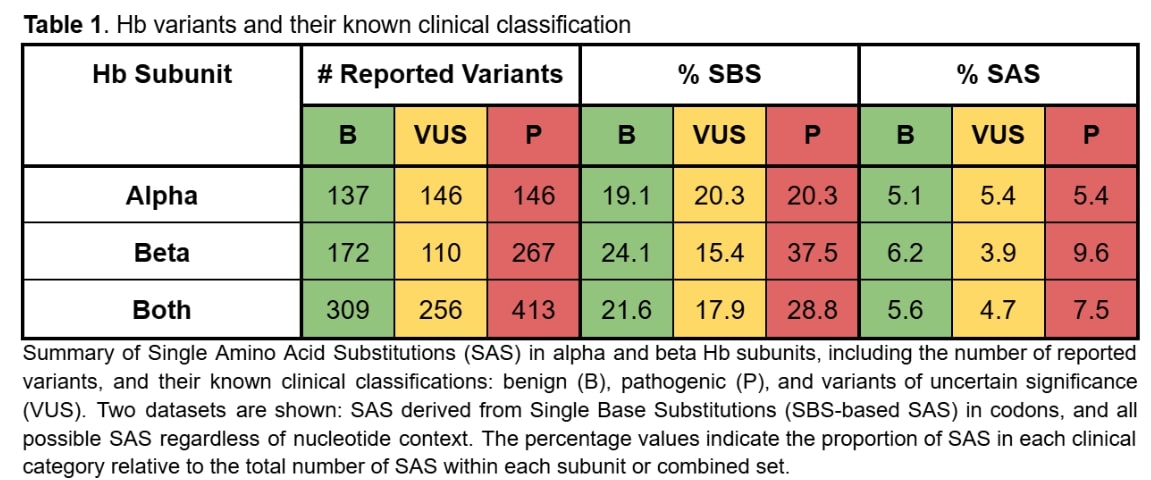
